# Changes in excitability and ion channel expression in neurons of the major pelvic ganglion in female type II diabetic mice

**DOI:** 10.1101/360826

**Authors:** Michael Gray, Kawasi M. Lett, Virginia B. Garcia, Cindy W. Kyi, Kathleen A. Pennington, Laura C. Schulz, David J. Schulz

## Abstract

Bladder cystopathy is a common urological complication of diabetes, and has been associated with changes in parasympathetic ganglionic transmission and some measures of neuronal excitability in male mice. To determine whether type II diabetes also impacts excitability of parasympathetic ganglionic neurons in females, we investigated neuronal excitability and firing properties, as well as underlying ion channel expression, in major pelvic ganglion (MPG) neurons in control, 10-week, and 21-week db/db mice. Type II diabetes in *Lepr^db/db^* animals caused a non-linear change in excitability and firing properties of MPG neurons. At 10 weeks, cells exhibited increased excitability as demonstrated by an increased likelihood of firing multiple spikes upon depolarization, decreased rebound spike latency, and overall narrower action potential half-widths as a result of increased depolarization and repolarization slopes. Conversely, at 21 weeks MPG neurons of db/db mice reversed these changes, with spiking patterns and action-potential properties largely returning to control levels. These changes are associated with numerous time-specific changes in calcium, sodium, and potassium channel subunit mRNA levels. However, Principal Components Analysis of channel expression patterns revealed that the rectification of excitability is not simply a return to control levels, but rather a distinct ion channel expression profile in 21-week db/db neurons. These data indicate that type II diabetes can impact the excitability of post-ganglionic, parasympathetic bladder-innervating neurons of female mice, and suggest that the non-linear progression of these properties with diabetes may be the result of compensatory changes in channel expression that act to rectify disrupted firing patterns of db/db MPG neurons.

## INTRODUCTION

Diabetes is a systemic, progressive disease characterized by lack of insulin directly (Type I), or functional lack of its effectors and/or insulin resistance (Type II), that ultimately leads to hyperglycemia and chronic complications. Diabetes is the seventh leading cause of death in the United States and afflicts 11.1-11.9% of adults over the age of 20; this rate has increased from 8.9% in years 1988-1994 to 11.9 % in years 2011-2014 (United States. Department of Health and Human Services. et al. 2016). In addition, diabetic complications are age dependent (Gunnarsson 1975; DCCT Research Group 1996; Liu et al. 2017). This age dependence combined with the 111.5% increase in persons age 65 and older from 1975-2015 (United States. Department of Health and Human Services. et al. 2016), necessitates studying an important complication of diabetes—diabetic neuropathy.

Diabetic neuropathy is particularly dangerous due to its insidious development, susceptibility of autonomic neurons and their subsequent regulation of vital organ systems, and a potential for positive feedback loops through dysregulation of microvasculature (Faerman et al. 1971; Vinik et al. 2003). Interventions are difficult as symptoms are subclinical and are often undiagnosed until long after neural lesions occur. That is, autonomic motor neuron damage occurs long before patients report somatic sensory symptoms. This is supported by the finding that in diabetic mice, thin, non-myelinated post ganglionic autonomic motor neurons show deficits long before somatic sensory motorneurons do (Liu et al. 2017). Neuropathic lesions can affect various autonomic pathways resulting in gastroparesis (Vinik et al. 2003), cardiovascular dysregulation (Vinik et al. 2003, 2011), sudomotor dysfunction (Liu et al. 2017), diabetic cystopathy (Faerman et al. 1971; Frimodt-Moller 1980; Kaplan et al. 1995; Vinik et al. 2003), and dysregulation of blood flow in the periphery (UKPDS 1998). Bladder dysfunction as a result of diabetes (i.e. diabetic cystopathy) is characterized by reduced bladder sensation, increased post-residual void volume and overactive bladder (Kaplan et al. 1995; Yuan et al. 2015), and was first formally described in diabetics in 1864 (Faerman et al. 1971). Some of the effects of diabetic cystopathy can be attributed to damaged bladder afferents. For example, it has been shown in streptozocin (STZ) treated rats (type I diabetes model) that diabetic cystopathy is correlated with reduced levels of nerve growth factor (NGF) in dorsal root ganglion neurons (Sasaki et al. 2002). Furthermore, when NGF is expressed at the bladder wall, the development of diabetic cystopathy and subsequent increased post residual void volume can be mitigated (Sasaki et al. 2004). However, little is known as to how diabetes impacts autonomic neurons innervating the bladder, and how this contributes to autonomic neuropathy and bladder cystopathy.

The major pelvic ganglion (MPG) of the mouse is the primary motor innervation of the urinary bladder. When examining motor neurons of this network in males, Tompkins et al. (2013), found that in post-ganglionic, parasympathetic, MPG neurons in Lepr^db/db^ mice had an enhanced number of excitatory post-synaptic potentials (EPSP) up to 20 seconds after pelvic nerve stimulation relative to control. Simultaneously, the authors observed no change in EPSP amplitude. However, STZ-treated mice did not differ in EPSP number and had significantly reduced amplitudes relative to control mice. This suggests that the type I and type II models are distinct in how they affect physiology of MPG neurons.

Furthermore, change in action potential properties is similarly dependent on diabetic type: after-hyperpolarization (AHP) duration was significantly decreased in Lepr^db/db^ mice, while being significantly increased in STZ mice (Tompkins et al. 2013). Finally, MPG neuron input resistance and resting membrane potential are decreased and depolarized, respectively, in Lepr^db/db^ mice but unaffected in STZ mice (Tompkins et al. 2013). These results suggest that both EPSP and intrinsic properties of MPG neurons are differentially modulated by the nature of the diabetic model.

In this study, we exploit the Lepr^db/db^ C57BL/6J mouse, to extend our understanding of how type II diabetes affects progression of changes in neuronal excitability in neurons of the MPG of female mice. Specifically, we examine how these conditions change neurons at 10- and 21-week time points by combining current clamp recordings of neuronal output with ion channel expression analyses in the MPGs of female mice. We predicted that firing properties and intrinsic properties should resemble that observed by Tompkins et al. (2013); with a general increase in excitability at week 21 to evoked firing rate, an increased RMP, decreased input resistance, and increased AHP duration. However, we observed a distinct effect: excitability of MPG neurons was initially increased at 10-weeks, but subsequently became less excitable again – towards baseline levels – at 21-weeks. These results suggest a compensatory change in MPG neurons in diabetic animals. We then went on to investigate potential underlying mechanisms for these changes via measuremnts of ion channel mRNA levels in MPGs from 10-week and 21-week diabetic females.

## METHODS

### Type II Diabetes Model – Lepr^db/db^ Animals

Wild type (WT) C57BL/6J and Lepr^db/db^ female mice (*Mus musculus*) were obtained from the Jackson Laboratory (Bar Harbor, ME, USA). Lepr^db/db^ mice have a mutation that interrupts the longest isoform of the leptin receptor (Chen et al. 1996). These animals display hyperphagia, obesity, hyperglycemia and hyperinsulinemia, which has led to their widespread use as a model of type II diabetes. Together, mouse and rat leptin mutants have been used in over 4000 published studies of type II diabetes (Wang et al. 2014). Lepr^db/db^ mice have been used by the Animal Models of Diabetic Complications Consortium and others as models for diabetic neuropathy (Sullivan et al. 2007).

Mice were group housed in cages on a 12 hr. light/dark cycle and fed a standard chow diet ad libitum. All animal procedures were performed in accordance with National Institutes of Health Guide for the Care and Use of Laboratory Animals and approved by the University of Missouri Animal Care and Use Committee.

### Establishing a Diabetic Phenotype

#### Glucose Measurement

At either 10 (n = 15 WT and 14 Lepr^db/db^) or 21 (n = 6 WT and 14 Lepr^db/db^) weeks of age animals were fasted for 4 hours and then a fasting glucose measurement was taken via tail blood using the average of two readings of a OneTouch^®^ (Sunnyvale, CA) or ReliOn Prime^®^ (Arkay inc., Kyoto, Japan; Distributed by Walmart, Bentonville, AR) Blood Glucose Monitoring System. At this time blood was collected to measure fasting serum insulin levels. Weights were also recorded at this time.

#### Insulin Elisa

Serum insulin levels were analyzed using a Mouse/Rat Insulin Elisa (Millipore^™^, Billerica, MA) according to manufacturer’s instructions. A total number of 6 samples at 10 weeks were run for both WT and Lepr^db/db^ females. At 21 weeks of age 9 Lepr^db/db^ and 6 WT females were analyzed. For Lepr^db/db^ serum was diluted at 1:20 with matrix solution provided in the ELSIA kit to allow for measurements to fall on the standard curve.

### Electrophysiology

MPGs were dissected from isoflurane euthanized female mice from wildtypes (n = 9) at 10 weeks (WT), Lepr^db/db^ at 10 weeks (n = 5; DB10), or Lepr^db/db^ at 21 weeks (n = 5; DB21). Dissection and all experiments were done in oxygenated physiological saline at room temperature. MPGs were dissected out and pinned to a Sylgard (Dow, Midland, MI) lined perfusion chamber. The preparation was then desheathed using small pins to remove excess tissue. Saline was composed of in mM: NaCl, 146; KCl, 4.7; MgSO_4_, 0.6; NaHCO_3_, 1.6; NaH_2_PO_4_, 0.13; CaCl_2_, 2.5; Glucose, 7.8; HEPES, 20; pH’d to 7.3. Electrodes were pulled on a P-97 microelectrode puller (Sutter, Novato, CA) filled with 3M KCl, and had resistances of 30 to 60 MΩ. Recordings were acquired in Bridge mode using an Axoclamp 900A and digitized using a Digidata 1440A using the pClamp 10.3 suite of software (Molecular Devices, Sunnyvale, Ca) running on an IBM-compatible computer. Silver electrode and ground wire were chlorided using household Bleach (Clorox, Oakland, CA).

#### Passive properties

Resistance, time constant, capacitance and rebound spikes were estimated from −500 pA current injections of 400 ms. In some protocols where there were no rebound spikes, these properties were estimated from −500 pA current injections for 2000 ms, To ensure these protocols did not differ, we examined action potential properties for 12 WT neurons where both protocols were used. A paired t-test showed that hyperpolarization duration did not significantly affect these measurements, the closest measurement to being affected was AHP (t (11) = 2.090, p = 0.061) while all other properties were unaffected (p > 0.1). Due to the contamination of slow activating currents and membranes being non-isopotential in real neurons, time was estimated by the 2 exponential fit; 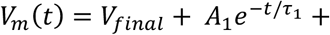 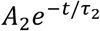; the larger A (IR) and time constant term were taken to be the time constant (Golowasch et al. 2009; White and Hooper 2013). Capacitance was estimated from this time constant and input resistance. For excitability data, current was injected from 0 to 700 pA in 50 pA steps for 400 ms.

#### Action Potential Properties

Action potential properties were estimated both manually (Spike count, Rheobase, AP slope to depolarizing current injection, and Threshold) and by using the auto-statistics package of the Clampfit program from the pClamp 10.3 suite of software (Molecular Devices, Sunnyvale, Ca) (All others). These properties are illustrated in figure 1 for reference, and unless otherwise stated were performed on rebound spikes to prevent contamination from current injection.

**Figure 1.**
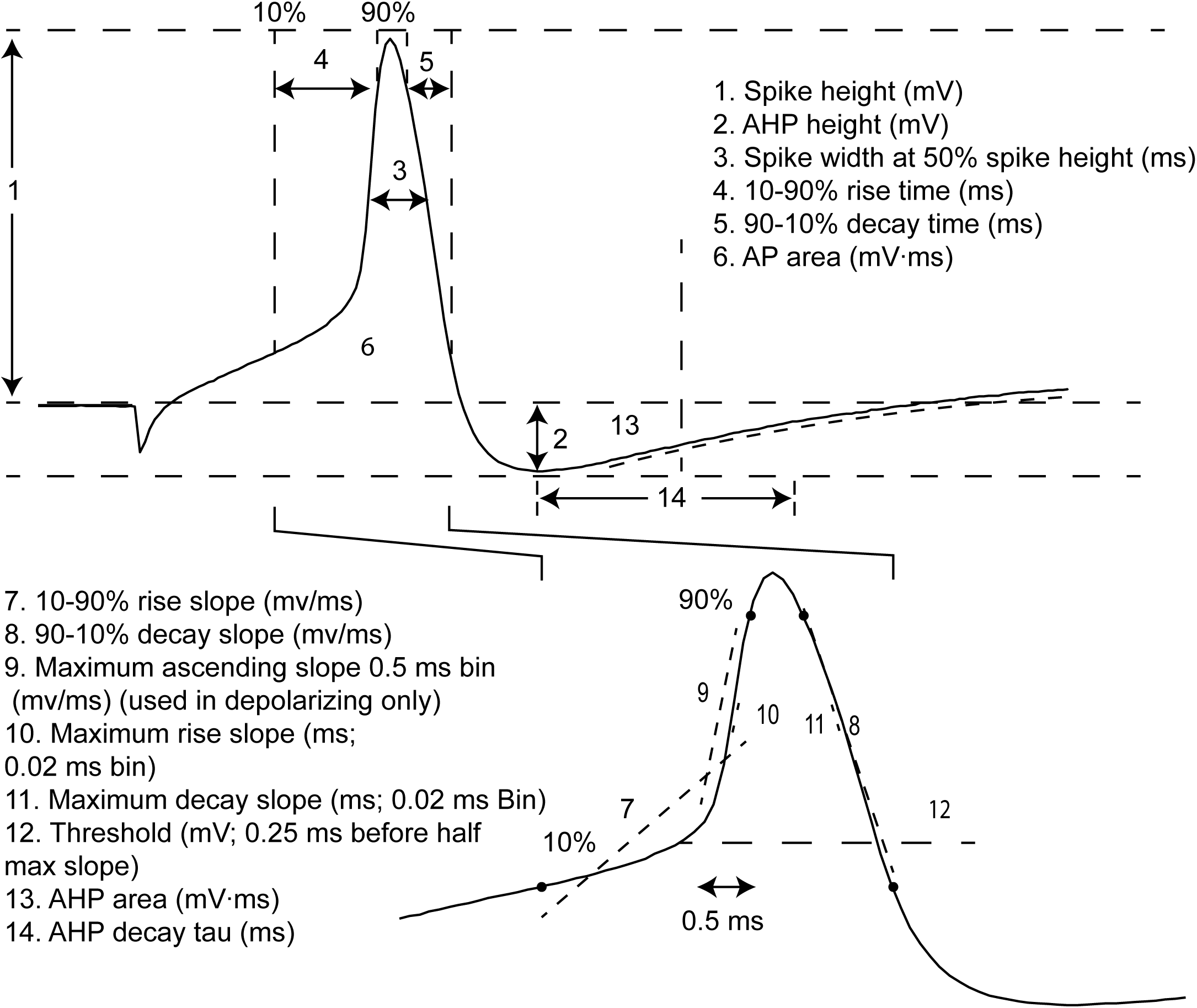
Action potential properties measured from depolarization and rebound induced spikes. Each number corresponds to a given measurement, as described in the Methods. These numbers are referred to in the Results section when relevant.

#### Threshold measurement

Threshold was measured from two methods and verified by estimation from passive properties and rheobase. First, we extrapolated manually from step protocols. Second, we adopted an adapted version of the spike slope method, where the derivative of voltage with respect to time slope is plotted vs voltage and the voltage at which this slope meets or exceeds some slope is threshold usually set from 2 to 20 mV/ms (Naundorf et al. 2006; Platkiewicz and Brette 2010, 2011). However, rather than using constant ascending spike slope, this method was adapted by measuring maximal ascending slope within a 0.5 ms bin, defining threshold as the voltage recorded 0.25 ms before the center point of the half maximal bin. Within wildtype, no significant differences were found between this method (M = −23.5, SD = 8.1) and manual estimation (M = −23.5, SD = 8.1). Slope method was chosen to minimize any confounds between changes in ascending slope.

#### Inclusion Criterion

Some neurons impaled produced no spikes. As we could not ascertain whether this was due to impalement damage, or these cells being silent neurons or closely apposed satellite glial cells (Hanani 2010), the following inclusion criteria were made: 1) All neurons must produce spikes to depolarizing current injections. 2) All cells must have resting membrane potentials less than −30 mV. 3) All cells must have input resistances greater than ~18 MΩ.

#### Statistics

Data was organized and stored in Microsoft Excel (Microsoft, Redmond, WA). All Statistics and graphs were made in Sigmaplot 11.0 (Chicago, Il), R (https://www.r-project.org/) and formatted in Adobe Illustrator CC 2017 (San Jose, CA). Data that passed normality testing and were shown to be homoscedastic were analyzed using one-way ANOVA. Most Data, however, was found to be non-normal and /or fail equal variance and comparisons were therefore made with a non-parametric Kruskal-Wallis one-way ANOVA on ranks using a post-hoc Dunn’s test. The two-way ANOVA for Figure 3B failed the assumptions of normality and homoscedasticity and could not be successfully transformed, statistics are reported for raw data.

### qRT-PCR Methods

Paired MPGs from each animal were collected into Trizol reagent (Life Technologies, Carlsbad, CA), homogenized, and stored at −80°C until RNA extraction. Total RNA was isolated from MPGs according to the protocol provided by the manufacturer. Complementary DNA (cDNA) was generated from 100 ng total RNA primed by a mixture of random hexamers and oligo-dT primers. Reverse transcription reactions were carried out at a volume of 20 μl using qScript cDNA SuperMix (QuantaBio, Beverly, MA) according to manufacturer protocols. Following cDNA synthesis, the reaction was heat inactivated and diluted 5X to a final volume of 100 μl before being used as a template for quantitative PCR (qPCR). From each cDNA reaction we quantified at least 15 different gene products. Multiple cDNA synthesis reactions were carried out from a single total RNA sample to quantify the full suite of genes examined in this study.

In our previous work, we designed or modified and independently validated qPCR primers for use in absolute quantitation of mRNA copy number for all of the genes of interest in this study. These primer sets are previously published, and standard curves generated and used as described in our previous work (Garcia et al. 2014, 2018). Briefly, qPCR reactions were carried out using SYBR mastermix (BioRad) according to the manufacturer’s instructions, and consisted of primers at final concentrations of 2.5 μM. Reactions were carried out on a CFXConnect (BioRad) machine with a three-step cycle of 95°C-15s, 58°C-20s, 72°C-20s, followed by a melt curve from 65°C to 95°C. Fluorescence data acquisition was made at the 72°C step, and every 0.5°C of the melt curve. All reactions were run in triplicate, and the average Ct (cycle threshold) was used for interpolation with standard curves to generate copy number for a given reaction.

The unit we use to express all of the qPCR data in this study is “copy number per ng of total RNA,” and reflects the amount of input RNA that went into the cDNA synthesis reaction. All of the data were normalized (see Garcia et al. 2014) relative to the average expression of Glyceraldehyde 3-phosphate dehydrogenase (GAPDH), beta-actin, and hypoxanthine guanine phosphoribosyl transferase (HPRT) genes from each sample (Vandesompele et al. 2002). Samples that were found to have low expression of these control genes were eliminated from the analysis. None of the control genes showed significant differences in expression across groups.

## RESULTS

### Lepr^db/db^ mice produce a diabetic phenotype with hyperglycemia that differs according to age

Diabetic neuropathy and the metabolic derangements of diabetes itself (Gunnarsson 1975; Giachetti 1978; Medici et al. 1999) are age-dependent. Therefore, we wanted to verify that our Lepr^db/db^ model exhibited diabetic physiological properties such as weight gain, elevated blood sugar and serum insulin to verify our diabetic model had sufficient time to develop the phenotype.

### Lepr^db/db^ mice are heavier than WT

Mean weights (Figure 2A) for diabetic mice from weeks 10 to 21, increased from 44.1 g (SD = 3.4) to 55.4 g (SD = 5.5), respectively (Holm-Šídák, p < 0.001) and were significantly heavier than WT at both time points (Holm-Šídák, p < 0.001). In contrast, there was not a statistically significant difference in weight for WT mice from week 10 (M = 18.9, SD = 1.6, g) to 21 (M = 20.0, SD = 1.2, g) (Figure 2A, Holm-Šídák, p = 0.082). [Two way ANOVA: Genotype; F (1, 43) = 169.319, p < 10^−8^. Age; F (1, 43) = 21.285, p = 3.5 x 10^−5^. Interaction; F (1, 43) = 3.401, p = 0.072].

**Figure 2.**
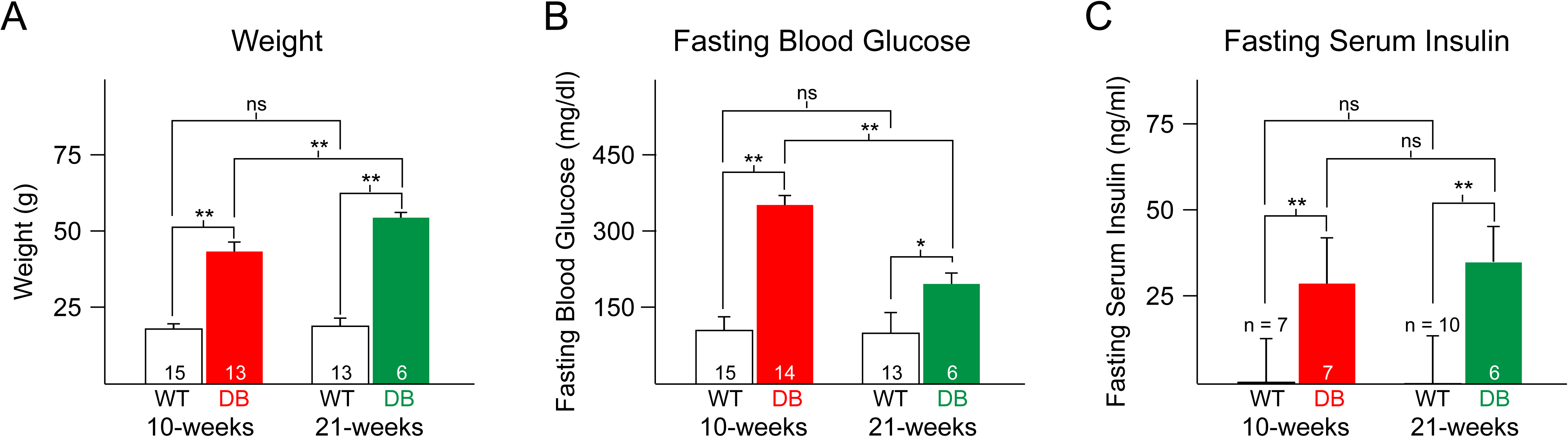
Lepr^db/db^ mice show a strong diabetic phenotype at week 10 with hyperglycemia that is somewhat reduced at week 21. Diabetes associated metabolic parameters of DB vs WT mice at different times; week 10 *(left)* or week 21 *(right)* in either wildtype (white) or diabetic conditions (red and green for weeks 10 and week 21 respectively). **A.** A two-way ANOVA for factors genotype and age showed that mouse weight was influenced by genotype and age. [Genotype; F (1, 43) = 169.319, p < 10^−8^. Age; F (1, 43) = 21.285, p = 3.5 x 10^−5^. Interaction; F (1, 43) = 3.401, p = 0.072] **B.** A two-way ANOVA for factors genotype and age showed that fasting blood glucose was influenced by genotype and age. [Genotype; F (1, 43) = 37.150, p = 2.7 x 10^−7^. Age; F (1, 43) = 8.893, p = 0.005. Interaction; F (1, 43) = 7.762, p = 0.008]. **C.** Two-way ANOVA for factors genotype and age showed that fasting serum insulin was influenced by genotype. [Genotype; F (1, 43) = 71.608, p < 10^−8^. Age; F (1, 43) = 0.0595, p = 0.809. Interaction; F (1, 43) = 0.0113, p = 0.916]. Holm-Sidak Post-Hoc test; *, p < 0.05; **, p < 0.01. Error bars are SEM.

### Lepr^db/db^ mice are hyperglycemic, especially when young

Surprisingly, in diabetic mice, fasting blood glucose (Figure 2B) significantly *decreased* from 352.54 mg/dl (SD = 115.22) at week 10 to 196.62 mg/dl (SD = 113.16) at week 21, this phenomena has been reported for C57BL/6J mice before, however, usually at later time points (~week 25-30; although still elevated compared to control (Gunnarsson 1975). Despite this, consistent with the diabetic phenotype, fasting blood glucose was significantly elevated in diabetic mice for both week 10 (Holm-Šídák, p < 0.001) and week 21 (HS, p = 0.044) vs WT (Figure 2B). In contrast to diabetic mice, no significant difference was observed in the fasting blood glucose of WT mice between weeks 10 (mg/dl, M = 112.47, SD = 27.14) and 21 (M = 107.17, SD = 8.72; Holm-Šídák, p = 0.901). [2-way ANOVA: Genotype; F (1, 43) = 37.150, p = 2.7 x 10^−7^. Age; F (1, 43) = 8.893, p = 0.005. Interaction; F (1, 43) = 7.762, p = 0.008].

### Lepr^db/db^ mice have elevated plasma insulin

In C57BL/6J Lepr^db/db^ mice the plasma insulin level is known to increase (Gunnarsson 1975). Mean plasma insulin levels (Figure 2C) in diabetic mice were stable from week 10 (ng/ml, M = 29.51, SD = 40.19) to week 21 (ng/ml, M = 35.11, SD = 46.51; Holm-Šídák, p = 0.908), but were significantly elevated compared to WT(Holm-Šídák, p < 0.001) which also did not change from weeks 10 (ng/ml, M = 0.51, SD = 0.43) to 21 (ng/ml, M = 0.33, SD = 0.07; Holm-Šídák, p = 0.817). Together, these results support that the Lepr^db/db^ manifests the insulin resistant, type II diabetic phenotype.

### Firing properties of Lepr^db/db^ neurons changes with diabetic condition and time

#### DB10 and DB21 neurons have enhanced and reduced excitability, respectively

Our hypothesis was that since diabetes causes diabetic cystopathy (Kebapci et al. 2007), presumably at least in part through diabetic autonomic dysregulation (Nadelhaft and Vera 1992), Lepr^db/db^ efferent bladder-innervating neurons of the MPG may contribute to these pathological changes and may manifest themselves as changes in intrinsic excitability. Therefore, we first examined whether different diabetic conditions would spike differently in response to depolarizing current steps from 0 to +700 pA in 50 pA increments (Figure 3A). We used 114 cells from 10-week-old WT mice (WT, n = 37); 5, 10-week-old diabetic mice (DB10, n = 41) and 5, 21-week-old diabetic mice (DB21, n = 36). As we saw no difference in metabolic parameters for WT mice from weeks 10 to 21 (Figure 2), we assume WT physiology is likewise similar to WT21. Thus all WT recordings are made from 10-week old animals. The representative neurons in Figure 3A show that DB10 (Red) neurons appear to be more excitable than WT (Black). In contrast, DB21 neurons (Green) were less excitable than either DB10 or DB21, and appeared to have reduced input resistance (discussed below). Surprisingly, across all conditions, of the 114 neurons studied, only 13 (11.4%) produced more than one spike at any current injection level. Of the neurons that spiked more than once, 76.9% (10/13) were in the DB10 condition compared to a 15.4% (2/13) and 7.7% (1/13) contribution from WT and DB21 groups respectively. The probability of finding a neuron spiking more than once was 5.4% (2/37), 24.4% (10/41) and 2.8% (1/36) for WT, DB10 and DB21 groups respectively. Supporting this difference in excitability, a Chi square test showed that these groups were not statistically independent (*X^2^* (2, N = 114) = 8.268, p = 0.016). Similarly; a Kruskal-Wallis showed this data was significantly different, however, this test is more dubious as all groups had median = 1(Table 1). Due to this low spike probability, we could not run more conventional IO curves. Therefore, instead of IO curves, we examined the cumulative probability of producing one or more spikes as a function of current injected. A two way ANOVA for factors current and condition showed that cumulative probability of firing 1 or more spikes as a function of injected current was reduced in DB21 condition [Condition; F (2, 1443) = 37.997 p < 1 x 10^−8^. Current; F (12, 1443) = 51.817, p < 1 x 10^−8^. Interaction; F (24, 1443) = 2.038, p = 0.002.]. The probability of a neuron firing at least once (Top, Figure 3C) was not significantly different between DB10 and WT neurons (Holm-Šídák, p = 0.883), in contrast, DB21 had reduced probability of firing at least one spike from current injections of 100 to 350 pA when compared to DB10 (Holm-Šídák p < 0.05) and from 100 to 400 pA when compared to WT (Holm-Šídák p < 0.05). Unfortunately, this particular distribution was neither normally distributed nor homoscedastic due to its binary nature. Supporting the reduced excitability of DB21 neurons was the finding that median rheobase (Left, Figure 3B) was increased in DB21 (Mdn = 300 pA, IQR = 450-200, pA) relative to WT (Mdn = 175, IQR = 262.5-100, pA) (Dunn’s test, p < 0.01) and DB10 (Mdn = 150, IQR = 250-100, pA) (Dunn’s test, p < 0.01) [Kruskal Wallis ANOVA on Ranks; H (2) = 16.181, p < 0.001]. Together, these data support DB21 mice having reduced MPG excitability.

**Figure 3.**
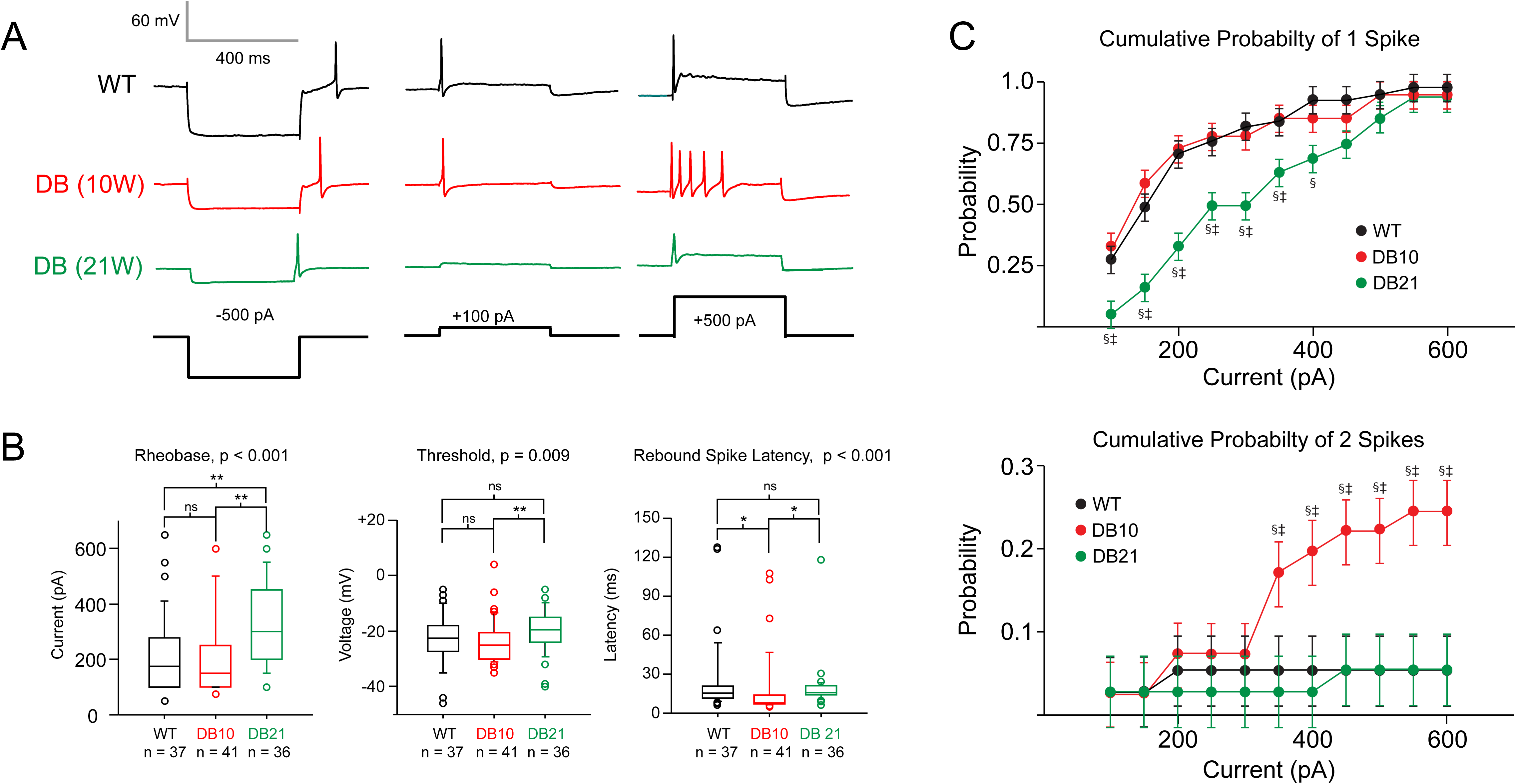
DB10 and DB21 conditions increase and decrease excitability of MPG neurons, respectively. **A.** Representative traces of intracellular currents injections of −500 pA, 100 pA and 500 pA current injections in WT10 (black), DB10 (red) or DB21 (green). **B.** *Left:* Kruskal-Wallis one way ANOVA on ranks showed that rheobase was significantly altered by diabetic condition [H (2) = 16.053, p < 0.001]. *Center:* A Kruskal-Wallis one way ANOVA on ranks showed that threshold (Figure 1, 12) was significantly altered by diabetic condition [H (2) = 9.351, p = 0.009]. *Right:* Kruskal-Wallis one way ANOVA on ranks showed that diabetic condition significantly altered latency to rebound spike [H (2) = 22.897, p < 0.001]. Shown are Medians, quartiles and outliers. Dunn’s test;*, p < 0.05; **, p < 0.01. ***, p < 0.001. **C.** *Top:* Two-way ANOVA for factors current and condition showed that cumulative probability of firing 1 or more spikes as a function of injected current was reduced in DB21 condition [Condition; F (2, 1443) = 37.997 p < 1 x 10^−8^. Current; F (12, 1443) = 51.817, p < 1 x 10^−8^. Interaction; F (24, 1443) = 2.038, p = 0.002.]. *Bottom:* A two-way ANOVA showed that the Cumulative probability of firing 2 or more spikes as a function of injected current was increased in the DB10 condition [Condition; F (2, 1443) = 43.428, p < 1 x 10^−8^. Current; F (12, 1443) = 2.251, p = 0.008. Interaction; F (24, 1443) = 1.301, p = 0.150.]. Note difference in scale. Holm-Sidak post hoc tests: p < 0.05; * WT10 vs DB10, † WT10 vs DB21,‡ DB10 vs DB21. Shown are Means ± SEM.

**Table 1.** Non-parametric (non-normal or non-homoscedastic) negative or redundant statistical summary † Internal control that is best fit line to sampled interval. ‡ n was 27, 32, 34 for WT, DB10 and DB21 respectively.

|  | WT<br>n = 35 |  |  | DB10<br>n = 34 |  |  | DB21<br>n = 34 |  |  | Kruskal-Wallis<br>p = |
| --- | --- | --- | --- | --- | --- | --- | --- | --- | --- | --- |
| Property (units) | Mdn. | 75<sup>th</sup><br>per. | 25<sup>th</sup><br>per. | Mdn. | 75<sup>th</sup><br>per. | 25<sup>th</sup><br>per. | Mdn. | 75<sup>th</sup><br>per. | 25<sup>th</sup><br>per. |  |
| 1<sup>st</sup> spike latency at depolarizing rheobase (ms) | 14.2 | 17.7 | 10.8 | 13.9 | 17.6 | 12.2 | 15.3 | 18.3 | 11.7 | 0.793 |
| 1<sup>st</sup> spike latency for Rebound spikes (-500 pA) | 15.2 | 22.3 | 11.6 | 7.9 | 13.9 | 7.9 | 15.7 | 21.2 | 14.1 | <0.001 |
| Baseline slope (mV/ms)<sup>†</sup> | -0.3 | -0.2 | -0.5 | -0.3 | -0.1 | -0.4 | -0.2 | -0.1 | -0.4 | 0.424 |
| Time from spike peak to AHP peak (ms) | 8.3 | 11 | 7.3 | 6.9 | 7.5 | 6.1 | 9.8 | 11.2 | 8.1 | <0.001 |
| Spike Height (mV) | 64.5 | 80.1 | 52.7 | 63.6 | 78.1 | 37.9 | 61.2 | 75.9 | 53.8 | 0.779 |
| Max Spikes (spikes) | 1.0 | 1.0 | 1.0 | 1.0 | 1.5 | 1.0 | 1.0 | 1.0 | 1.0 | 0.004 |
| Rebound spikes (-500 pA; spikes) | 1.0 | 1.0 | 1.0 | 1.0 | 1.0 | 1.0 | 1.0 | 1.0 | 1.0 | 0.628 |
| Change in Membrane potential 145 ms after AP repolarization (mV) ‡ | 1.8 | 3.8 | 0.9 | -0.2 | 2.0 | -0.7 | 0.8 | 1.8 | 0.2 | 0.005 |

**Table 2.** Negative or redundant statistical summary that was normal and homoscedastic. † Internal control that is essentially the RMP right after rebound. ‡ Redundant with area 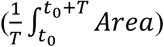

|  | WT | DB10 | DB21 | One- way<br>ANOVA<br>p = |
| --- | --- | --- | --- | --- |
| Property (units) | Mean (SD)<br>n = 35 | Mean (SD)<br>n = 34 | Mean (SD)<br>n = 34 |  |
| AP Mean Height, (mV) ‡ | 1.8 (2.5) | 0.7(1.7) | 1.7(1.6) | 0.045 |
| Baseline (mV)<sup>†</sup> | -41.0 (7.8) | -41(7.6) | -38.8(5.9) | 0.378 |

#### DB10 condition enhances excitability

When examining the probability of these neurons firing twice or more, neurons from DB10 had significantly greater probability of firing more than two spikes compared to both WT and DB21 (Bottom, Figure 3C). This change was significant from current ranges 350-700 pA vs DB21 (Holm-Šídák, p < 0.05) and 400-700 pA vs WT (Holm-Šídák, p < 0.05) [Condition; F (2, 1443) = 43.428, p < 1 x 10^−8^. Current; F (12, 1443) = 2.251, p = 0.008. Interaction; F (24, 1443) = 1.301, p = 0.150.]. Median DB10 rheobase (Left, Figure 3B; Mdn 150, IQR = 250-100, pA) was not significantly different from WT (Mdn 150, IQR = 275-100, pA) (Dunn’s, p > 0.05), but was significantly less compared to DB21 (Mdn 300, IQR = 450-200, pA) (Dunn’s, p < 0.05) [Kruskal-Wallis, H (2) = 16.053, p < 0.001]. In agreement with this, DB10 threshold (Center, Figure 3B; Mdn = −25.0, IQR = −20.5-(−30), mV) was hyperpolarized compared to DB21 (Mdn = −19.5, IQR = −15-(−24), mV) (Dunn’s test, p < 0.01), however, it was not significantly different from WT (Mdn = −22.0, IQR = −18-(−27.5), mV) but [Kruskal Wallis ANOVA on Ranks; H (2) = 9.351, p =0.009]. These results, suggest that DB10 excitability is enhanced relative to WT and DB21 conditions.

#### Rebound spike latency is decreased in DB10 neurons

Another sign that excitability was increased in DB10 neurons was that latency to first spike after release of hyperpolarizing current injection (Right, Figure 3B) was significantly decreased in the DB10 condition (; Mdn = 7.9, IQR = 13.9-7.1, ms) compared to WT (Mdn = 15.4, IQR = 20.9-11.5, ms)(HS, p < 0.05) and DB21 (Mdn = 15.7, IQR = 21.2-14.1, ms) (Holm-Šídák, p < 0.05) conditions. These data support excitability being enhanced in the DB10 condition.

### Passive properties of diabetic neurons change with age and diabetic condition

In order to characterize the mechanism of observed changes in firing of MPG neurons, we examined passive properties with the prediction that we should observe depolarized resting membrane potential (RMP) and reduced input resistance in diabetic neurons. In contrast to Tompkins et al. (2013), our data showed no difference in median resting membrane potential (Figure 4A) between WT (Mdn = −43.0, IQR = −36.5-(−47.0), mV), DB10 (Mdn = −42.0, IQR = −39.5-(−48.5), mV) or DB21 (Mdn = −40.0, IQR = −38.0-(−45.5), mV) (Figure 4A) [Kruskal-Wallace ANOVA on ranks; H (2) = 2.018, p =0.365]. However, consistent with Tompkins et al.and our finding of rheobase being increased in DB21, we found that median input resistance (Figure 4B) was significantly reduced in DB21 (Mdn = 42.5, IQR = 51.8-37.3, MΩ) (Dunn’s test; p < 0.004) but not in DB10 (Mdn = 51.0, IQR = 89.0-38.0, MΩ) (Dunn’s test, p > 0.05) neurons vs WT (Mdn = 68, IQR = 89.0-44.0, MΩ) neurons [Kruskal-Wallis ANOVA on ranks, H (2) = 10.641, p =0.005]. We next estimated time constant by a 2-exponential fit (Golowasch et al. 2009; White and Hooper 2013). Median time constant (Figure 4C) showed a similar pattern, showing a significant reduction in DB21 (Mdn = 1.8, IQR = 2.5-1.5, ms) compared to WT (Mdn = 2.3, IQR = 3.2-1.6, ms) (Dunn’s test, p < 0.01), but not compared to DB10 (Mdn = 2.5, IQR = 4.2-1.7, ms)(Dunn’s test, p > 0.05 [Kruskal-Wallis one way ANOVA on ranks, H (2) = 6.805, p =0.033]. Capacitance (Figure 4D) showed a different pattern, where both DB10 (Mdn = 48.7, IQR = 57.8-39.6, pF) and DB21 (Mdn = 41.9, IQR = 54.8-35.1, pF) were significantly increased relative to WT (Mdn = 32.9, IQR = 40.2-29.8, pF) (Dunn’s test, p < 0.01) [[Kruskal-Wallis one way ANOVA on ranks, H (2) = 23.187, p < 0.001]. These data suggest that capacitance is increased with diabetic condition independent of time, while time constant and resistance change with diabetic condition and age.

**Figure 4.**
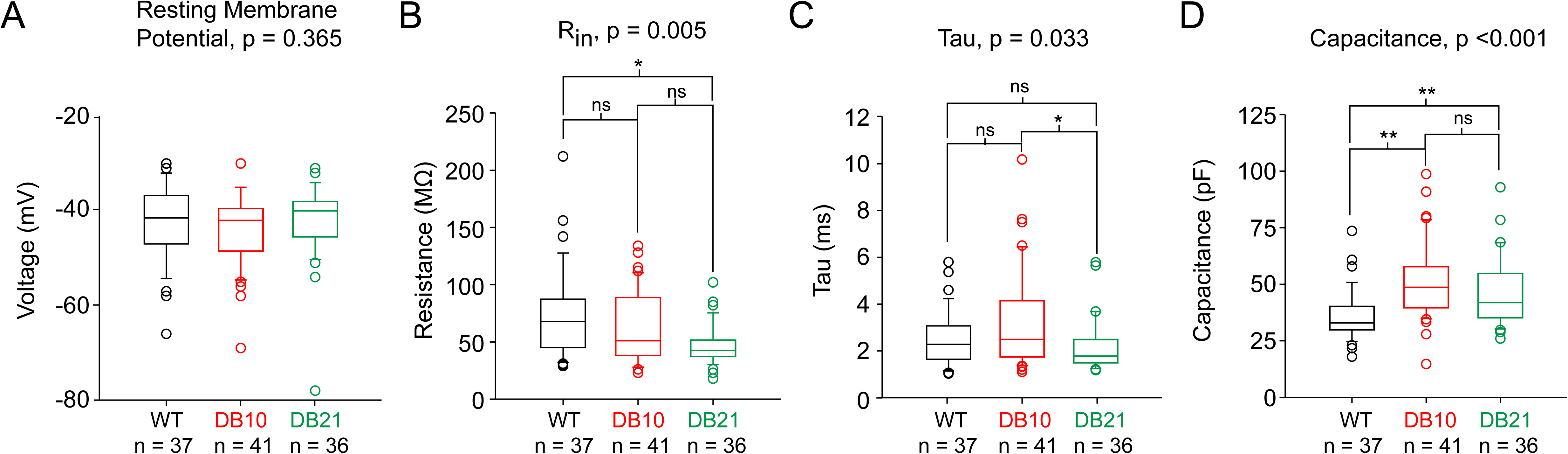
Effect of condition for WT, DB10, and DB21 condition on passive properties of MPG neurons. **A.** A Kruskal-Wallis one-way ANOVA on ranks showed that diabetic condition did not affect resting membrane potential (RMP) [H (2) = 2.017, p =0.365]. **B.** A Kruskal-Wallis one-way ANOVA on ranks showed that diabetic condition altered input resistance (R_in_) [H (2) = 10.641, p =0.005] **C.** A Kruskal-Wallis one way ANOVA on ranks showed that diabetic condition altered membrane time constant [H (2) = 6.805, p =0.033] **D.** A Kruskal-Wallis one-way ANOVA on ranks showed that diabetic condition altered capacitance of MPG neurons [H (2) = 23.187, p < 0.001]. Data shown are medians, quartiles and outliers. Dunn’s test: * p < 0.05; **, p < 0.01.

### Action potential properties in diabetic neurons change with age

In order to establish how observed changes in excitability caused by diabetic condition occurred, we examined whether diabetic condition and age modulated properties of the action potentials shown in figure 1.

#### Ascending spike slope is increased in response to depolarizing current injection in the DB10 condition dv but not first rebound spike

We looked at ascending action potential slope 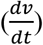 in response to depolarizing current injections (Figure 1, 9). DB10 mice had significantly greater median ascending spike slope (Figure 5A, Mdn = 30.0, IQR = 52.5-9.5, mv/ms) compared to either WT (Mdn = 8, IQR = 19.0-3.3, mv/ms) (Dunn’s Test; p < 0.01) or DB21 mice (Mdn = 5, IQR = 20.3-3.0, mv/ms) (Dunn’s Test; p < 0.01) [Kruskal-Wallis ANOVA on Ranks; H (2) = 22.365, p < 0.001]. Interestingly, in contrast to evoked spikes, when we examined median maximum ascending slope (i.e. bins of 0.02 ms; Figure 1, 10), in rebound spikes, DB10 neurons (Figure 5C, Mdn = 62.8, IQR = 77.9-40.3, mv/ms) did not significantly differ from WT (Mdn = 55.3, IQR = 77.9-40.3, mv/ms) or DB21 (Mdn = 40.3, IQR = 56.6-25.6, mv/ms)[Kruskal-Wallis ANOVA on Ranks; H (2) = 5.670, p = 0.059]. We believe this difference was real and not due to a reduction in statistical power, as within WT groups, the coefficient of variance in ascending slope for depolarizing current injections was greater (CV = 1.14) than for rebound spikes (CV = 0.79). In support of there being no differences in rebound spikes, when ascending spike slope was quantified using a 10-90% rise slope criterion (Figure 1, 7), DB10 neurons (Figure 5G, Mdn = 29.8, IQR = 40.6-15.4, mv/ms) did not significantly differ from either WT (Mdn = 25.8, IQR = 39.5-12.4, mv/ms) or DB21 (Mdn = 20.0, IQR = 32.7-11.5, mv/ms) [Kruskal-Wallis ANOVA on Ranks; H (2) = 2.043, p = 0.360]. These data support spike slope increasing in the DB10 condition in response to depolarization driven spikes but not rebound spikes.

**Figure 5.**
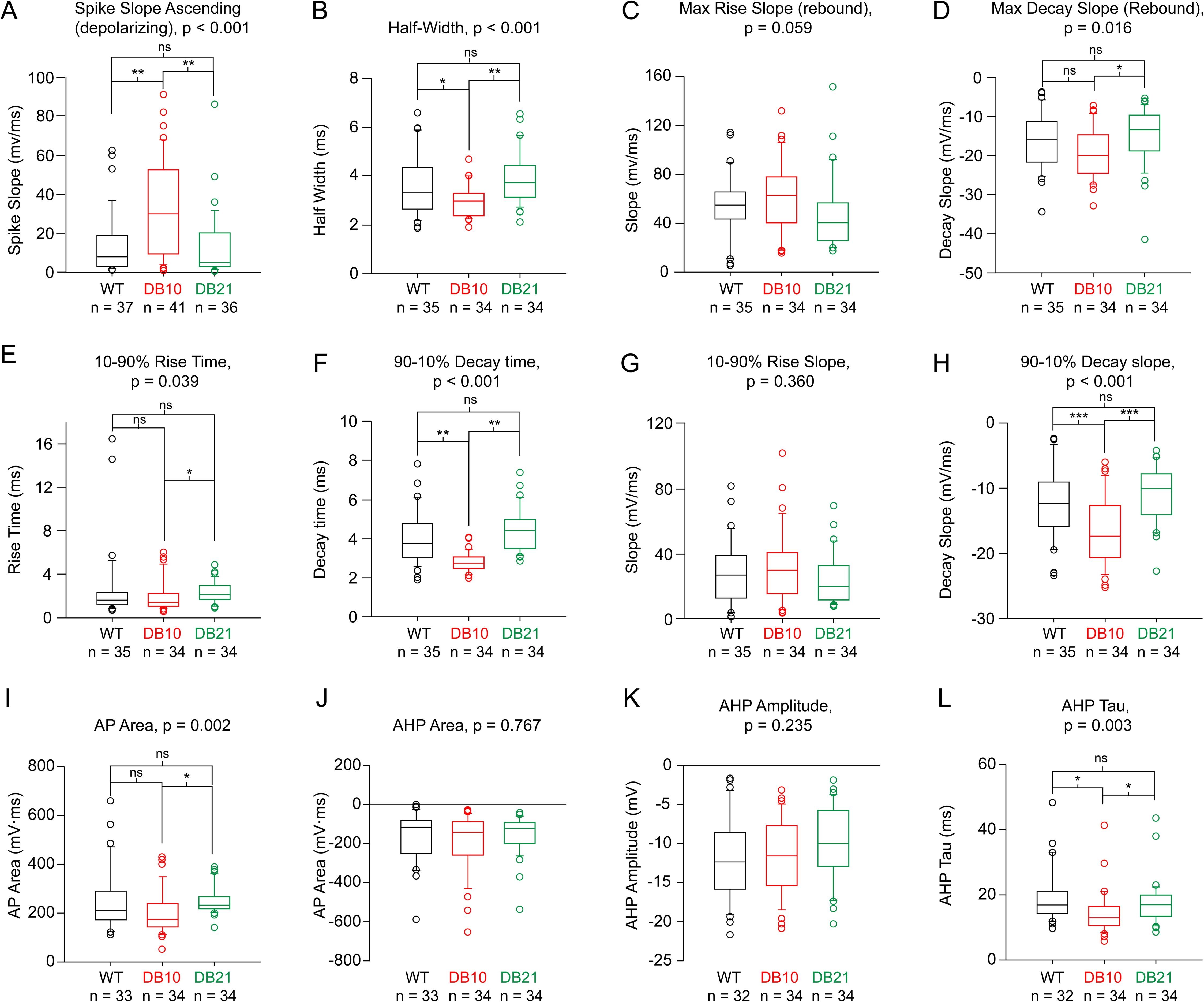
Action potential (AP) parameters are altered by DB condition in mouse MPG neurons. AP properties illustrated in figure 1 were quantified in 10 week wildtype (Black), 10 week Diabetic (Red) and 21 week diabetic (Green) in mouse MPG neurons. All data except A were quantified from rebound spikes. **A.** A Kruskal-Wallis one way ANOVA on ranks showed that diabetic condition affected ascending spike slope elicited by depolarizing current injection (Figure 1, 9) during depolarizing current injection of MPG neurons [H (2) = 22.010, p < 0.001]. **B.** A Kruskal-Wallis one way ANOVA on ranks showed that diabetic condition affected AP half-width (Figure 1, 3) [H (2) = 14.892, p < 0.001]. **C.** A Kruskal-Wallis one way ANOVA on ranks showed that diabetic condition did not significantly alter AP maximum rise slope (Figure 1, 10) [H (2) = 5.670, p = 0.059]. **D.** A Kruskal-Wallis one way ANOVA on ranks showed that diabetic condition altered maximum AP decay slope of (Figure 1, 11) [H (2) = 8.112, p = 0.017]. **E.** A Kruskal-Wallis one way ANOVA on ranks showed that diabetic condition altered AP 10-90% rise time (Figure 1, 4) [H (2) = 6.464, p = 0.039]. **F.** A Kruskal-Wallis one way ANOVA on ranks showed that diabetic condition altered 90-10% decay time (Figure 1, 5) [H (2) = 42.223, p < 0.001]. **G.** A Kruskal-Wallis one way ANOVA on ranks showed that diabetic condition did not alter AP 10-90% rise slope (Figure 1, 7) [H (2) = 2.043, p = 0.360]. **H.** A one-way ANOVA showed that diabetic condition altered 90-10% decay slope (Figure 1, 8) [F (2, 101) =11.541, p = 3.1 x 10^−5^]. **I.** A Kruskal-Wallis one-way ANOVA on ranks showed that diabetic condition altered AP area (Figure 1, 6) [H (2) = 12.320, p = 0.002]. **J.** A Kruskal-Wallis one-way ANOVA on ranks showed that diabetic condition did not alter AHP area (Figure 1, 13) [H (2) = 0.530, p = 0.767)]. **K.** A Kruskal-Wallis one-way ANOVA on ranks showed that diabetic condition did not alter AHP amplitude (Figure 1, 2) [H (2) = 2.900, p = 0.235)]. **L.** A Kruskal-Wallis one way ANOVA on ranks showed that diabetic condition altered AHP decay tau (Figure 1, 14) [H (2) = 11.896, p = 0.003)]. Data shown are medians, quartiles and outliers. Dunn’s test: * p < 0.05; **, p < 0.01.

#### AP half-width is decreased while decay slope is increased in rebound spikes of DB10 neurons

In contrast to ascending slope, we found that the median width of the action potential at half its maximal height (i.e. half-width; Figure 1, 3) was significantly reduced in the DB10 condition (Figure 5B, Mdn = 3.0 IQR = 3.3-2.4, ms) compared to DB21 (Mdn = 3.7, IQR = 4.4-3.1, ms) (Dunn’s test, p < 0.01) but not vs WT (Mdn = 3.3, IQR = 4.4-2.6, ms) (Dunn’s test, p > 0.05) [Kruskal-Wallis ANOVA on Ranks; H (2) = 14.991, p < 0.001].

Part of the reduction in DB10 half-width may be explained by the finding that that both maximal and 90-10% decay slope increased (Also see discussion). This is shown as median maximal decay slope (Figure 5D) was made more negative (increased) in DB10 neurons (Mdn = −20.0, IQR = −14.7-(−24.6), mv/ms) vs DB21 neurons (Mdn = −13.4, IQR = −9.7-(−18.9), mv/ms) (Dunn’s test, p < 0.05) but not vs WT (Mdn = −16.0, IQR = −11.3-(−22.1), mv/ms) (Dunn’s test, p > 0.05) neurons [Kruskal-Wallis ANOVA on Ranks; H (2) = 8.112, p = 0.017] suggesting these neurons repolarized faster. In agreement with this, mean 90-10% decay slope (Figure 5H) also became more negative in the DB10 condition (M = −16.7, SD = 5.3, mv/ms) vs WT (M = −12.4, SD = 5.6, mv/ms) (Holm Sidak, p = 0.001) and compared to DB21(M = −11.0, SD = 4.2, mv/ms), mv/ms)(Holm-Sidak, p < 0.001) [One way ANOVA; F (2, 100) =11.541, p = 3.1 x 10^−5^]. Accordingly, median 90-10% decay time (Figure 5F) significantly decreased in DB10 (Mdn = 2.7, IQR = 3.1-2.5, ms) vs WT (Mdn = 3.7, IQR = 4.8-3.0, ms) (Dunn’s test, p < 0.01) and DB21 (Mdn = 4.4, IQR = 5.0-3.5, ms) (Dunn’s test, p < 0.01) [Kruskal-Wallis ANOVA on Ranks; H (2) = 42.223, p < 0.001]. This data, in combination with the finding that AP ascending slope did not increase in rebound spikes, suggests that AP half-width decreased primarily through a mechanism that increases AP descending slope.

#### AP area and AHP decay constant decreased in DB10 with no changes in after hyperpolarization (AHP) area

Consistent with decreased AP half-width, total AP area (excluding the AHP; Figure 5I) was significantly decreased in DB10 neurons (Mdn = 174.4, IQR = 238.8-142.7, mV·ms) relative to DB21 (Mdn = 232.5, IQR = 266.7-217.5, mV·ms)(Dunn’s test, p < 0.05) but not when compared to WT (Mdn = 209.8, IQR = 290.7-172.0, mV·ms) (Dunn’s test, p > 0.05) [Kruskal-Wallis ANOVA on Ranks; H (2) = 12.320, p = 0.002]. This was not due to changes in spike height, as neither DB10 (Mdn = 63.6, IQR = 78.1-37.9, mV) nor DB21 (Mdn = 61.2, IQR = 75.9-53.8, mV) differed significantly in spike height (Table 1) relative to WT (Mdn = 64.5, IQR = 80.1-52.7, mV) [Kruskal-Wallis ANOVA on Ranks; H (2) = 0.498, p = 0.779]. If AP area decreased, a possible mechanism would be activation of outward currents, and one might expect to see an enhanced AHP. However, no change in AHP area (Figure 5J) for DB10 (Mdn = −141.6, IQR = −88.1-(−259.2), mV·ms), nor DB21 (Mdn = −121.9, IQR = −92.1-(200.5), mV·ms) vs WT (Mdn = 116.6, IQR = −80.6-(−251.1), mV·ms) was observed [H (2) = 0.530, p = 0.767)]. Neither were changes were observed in AHP height (figure 1, 2) in DB21 (Figure 5K, Mdn = −10.0, IQR = −5.8-(−12.9), mv), DB10 (Mdn = −11.6, IQR = −7.7-(−15.4), mv) conditions vs WT (Mdn = −12.4, IQR = −8.5-(−15.9), mv) [Kruskal-Wallis ANOVA on Ranks; H (2) = 2.900, p = 0.235]. Despite our finding of stable AHP amplitude and area, we did see that AHP decay tau significantly decreased in DB10 neurons (Mdn = 13.0, IQR = 16.5-10.5, ms) vs WT (Mdn = 16.9, IQR = 21.2-16.9, ms) (Dunn’s, p < 0.01) and DB21 (Mdn = 17.0, IQR = 20.0-13.4, ms) (Dunn’s, p < 0.05) (Figure 5L) [Kruskal-Wallis ANOVA on ranks; H (2) = 11.896, p = 0.003)]. These data suggest that diabetic condition and age affect interact to alter AP area and AHP decay constant while leaving other AP parameters unmodified.

#### 2^nd^ AHPs and ADPs after 145 ms

When quantifying properties of the AHP, it was found that after the initial AHP, some neurons had after depolarizations, while some had secondary slower AHPs and still others had both. We attempted to systematically quantify this across conditions by analyzing the change in voltage 145 ms after the descending stroke of the AP recrossed baseline. However, this comes with the caveat that some neurons could not be quantified due to no change in baseline, multiple rebound spikes, or loss of cells. Within 73.0 % (27/37) of quantified WT neurons, 81.5% (22/27) displayed afterdepolarizations, while the remaining 18.5% (5/27) displayed secondary after hyperpolarizations and none displayed both. Within DB10 neurons 4.9% (2/41) could not be classified due to multiple rebound spikes. However, of the 82.9% (34/41) that could be quantified, 44.1% (15/34) showed ADPs, 20.6% (7/34) showed secondary AHPs; in contrast to WT; where 29.4% (10/34) of these neurons showed both AHPs and ADPs. Within 94.4% (34/36) of quantified DB21 neurons, 76.5% (26/34) showed ADPs, 2.9% (1/34) showed secondary AHPs, while 20.6% (7/34) showed both. A Chi Square test showed that these groups were statistically independent (*X^2^* (4, N = 95) = 15.960, p = 0.003). Alternatively, when we just quantified the absolute voltage change at 145 ms, DB10 (Mdn = −0.2, IQR = 2.0-(−0.7), ΔmV) (Dunn’s, p < 0.05) but not DB21 (Mdn = 0.8, IQR = 1.8-0.9, ΔmV) (Dunn’s, p > 0.05) was significantly different from WT (Mdn = 1.8, IQR = 3.8-0.9, ΔmV) (Table 1) [Kruskal-Wallis ANOVA on Ranks; H (2) = 10.766, p = 0.005]. This data suggests that the DB10 condition is more likely to have slow AHPs or a lack of ADPs relative to WT.

### Ion Channels Show Widespread Changes in Expression in Both DB10 and DB21 Animals

We specifically focused on subsets of ion channel and receptor genes for this study to start to understand the underlying mechanisms for changes in excitability that we detected in MPG neurons of diabetic animals. We sampled alpha subunits for voltage-dependent Ca^2+^ channels *(CACNA1A-H)*, a subset of Na^+^ channels *(SCNx)*, and six families of K^+^ channels *(KCNA, KCNB, KCNC, KCND, KCNN, KCNMA)*. Of 30 ion channel genes studied, 8 were significantly different between DB10 and control wild-type animals (Figure 6). The voltage-dependent Ca^2+^ channels *CACNA1A, CACNA1E* and *CACNA1H* were significantly higher in DB10 animals relative to control. The only Na^+^ channel gene that was changed was a significant increase in *SCN2A1* in DB10 animals. In addition, 4 K^+^ channel genes showed altered expression in DB10 animals: *KCNA3* was significantly lower in DB10 animals, while *KCNB1, KCNC2*, and *KCND3* were significantly higher in DB10 mice.

**Figure 6.**
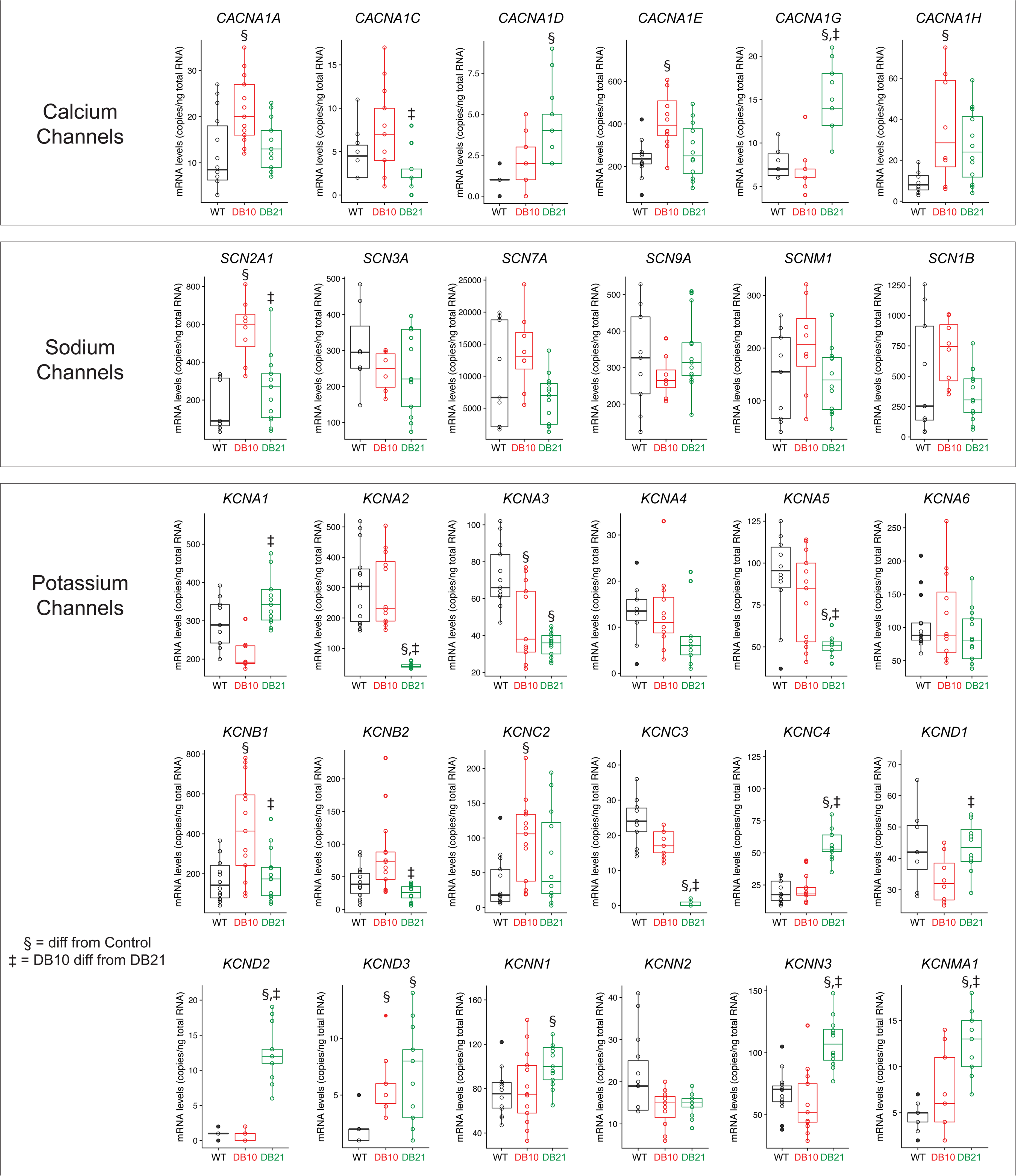
mRNA copy numbers for ion channel subunits of across WT, DB10, and DB21 experimental groups. Significant differences as noted (p < 0.05; post-hoc Holm-Šídák analyses following One-Way ANOVA) represent pairwise comparisons across all three groups. Data shown are medians, quartiles and each individual value from a given animal.

In DB21 animals, 12 channel genes showed differential expression relative to wild-type controls (Figure 6). *CACNA1D* and *CACNA1G* were significantly higher in DB21 animals relative to control. There were no significant differences in sodium channel gene expression seen between DB21 and wild-type animals. However, there were 10 different K^+^ channel genes that showed changes in expression in DB21 animals. *KCNA2, KCNA3, KCNA5*, and *KCNC3* all were significantly lower in DB21 animals than wild-type, with *KCNA2* and *KCNC3* expression all but abolished in the DB21 animals. *KCNC4, KCND2, KCND3, KCNN1, KCNN3* and *KCNMA1* were all significantly higher in DB21, with *KCND2* expression only detectable in DB21 animals. Most of these changes are seen only in DB21 animals and not in DB10, including *KCNA2, KCNA5 KCNC3, KCNC4, KCND2, KCNN1, KCNN3* and *KCNMA1*.

Some changes seen in DB10 animals resolve to control levels in the DB21 animals, while others persist throughout the DB21 time point (Figure 6). *KCNA3* is significantly lower in DB10 animals, and this change persists into the DB21 group. *KCND3* is significantly increased in both DB10 and DB21 animals. Conversely, *KCNB1* is significantly higher in the DB10 animals, but returns to control levels in DB21 animals. Finally, there are some channels that are significantly different only between DB10 and DB21. *KCNA1* and *KCND1* are significantly higher in DB21 than DB10, although there is a trend towards these two channels being downregulated in DB10 overall, even though this does not reach statistical significance relative to control. The converse is true for *KCNB2*, which is significantly higher in DB10 than DB21 – although there is a trend for this channel to be transiently upregulated in the DB10 group.

### Principal Components Analysis to Further Examine Differences Among Groups in Firing Properties and Channel Expression

We next used principal component analysis (PCA) to visualize potential patterns and correlations in the features that may underlie distinctions in both the spike properties and channel expression across groups. A variance plot of the first two principal components (PC) for spike characteristics (Figure 7A) demonstrates that there is overlapping features in all three groups that do not distinguish these groups in any obvious way. There is one major PC (PC1) that accounts for the majority of the variance in the data 42.5%, Figure 7B). The variables that most contribute to the variance in spike characteristics largely consist of features involved in shaping the action potential dynamics underlying spike shape, such as decay slope, rise slope, and the amplitude of the spike (e.g. spike height, peak amplitude). These would consistent with changes in Na^+^ and K^+^ channel expression, particularly those subtypes involved in the spiking itself. However, there is no combination of features that separates the diabetic animals from the wild-types, or each other, in a discrete fashion.

**Figure 7.**
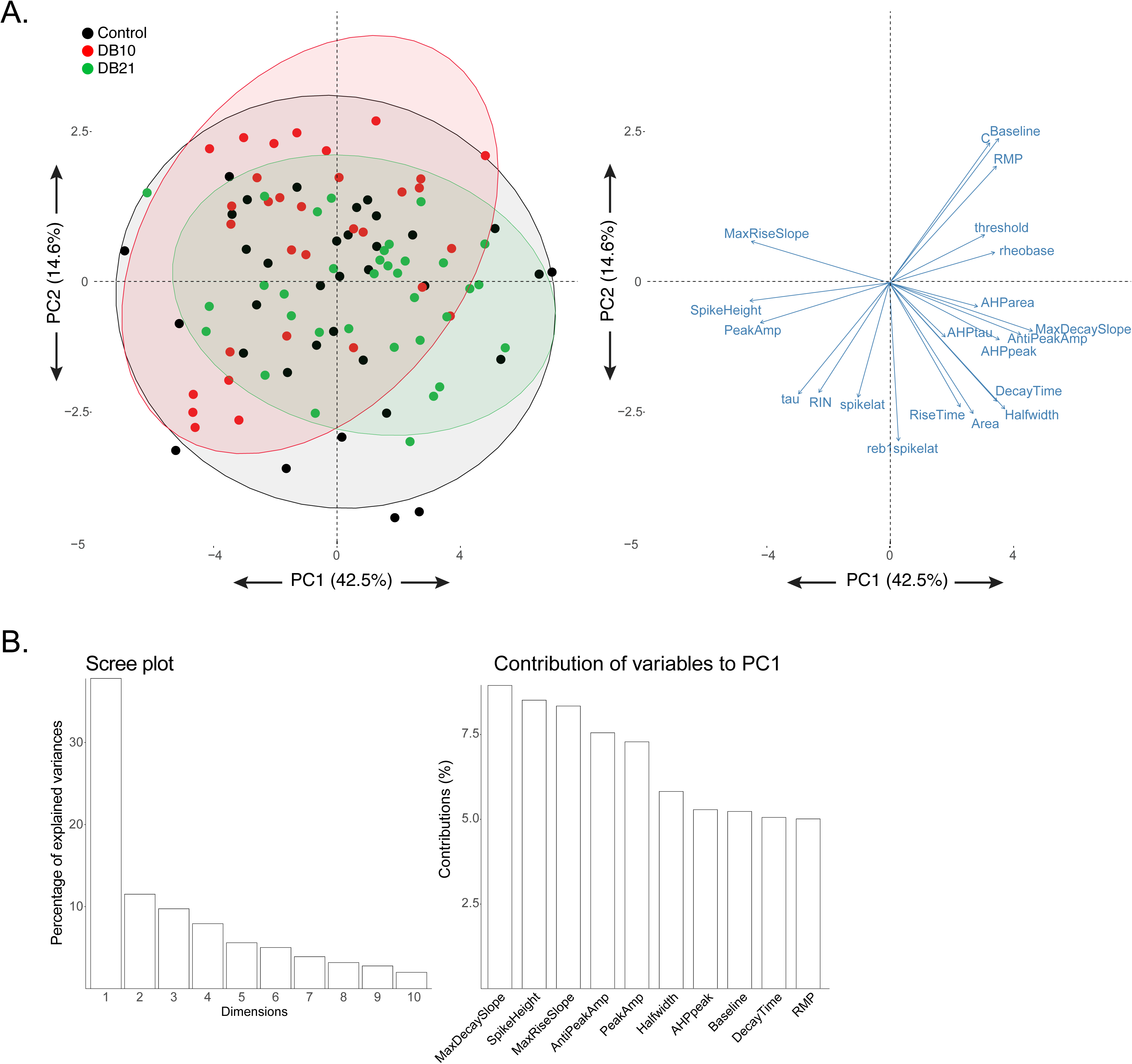
Principal components analysis (PCA) of spike characteristic across WT, DB10, and DB21 mice. **A.** The first two principal components (PC1 and PC2) define the x-and y-axes respectively. PC1 accounted for 42.5% of the variance while PC2 accounted for 14.6%. By and large, all three groups show overlapping distribution of variance across PC1 and PC2. **B.** Scree plot demonstrating the amount of variance accounted for across the first 10 principal components. A substantial plurality of the variance is accounted for in the first principal component, while the remainder contribute far less to the variance in the data. C. Post-hoc analysis of the variables that contribute to the variance in PC1 reveals that relatively equal contributions are found from variables associated with spike shape and AHP amplitude.

The PCA for channel expression in the MPG tells a different story. The variance plot of PC1 and PC2 (Figure 8A) shows clear separation of all three groups, and the scree plot (Figure 8B) notes that these two PCs account for the majority of the variance in the data. When looking at the channels that contribute to PC1 and PC2, it is also clear validation of the changes in expression reported in Figure 6. By examining the vector plot in Figure 8A and the contributions to PC1 in Figure 8B, it is possible to discriminate a group of channels largely responsible for distinguishing the DB21 animals. The top ten contributors to PC1 consists entirely of channels that are uniquely differentially expressed in DB21 animals relative to both DB10 and control. Conversely, by examining the top ten contributors to PC2 – in conjunction with the vector plot – it can be seen that PC2 consists of channels uniquely differentially expressed in DB10 animals *(CACNA1A, CACNA1E, SCN2A1, KCNB1, KCNC2)* as well as channels that are both changed in the same direction in DB10 and DB21 *(KCND3)*.

**Figure 8.**
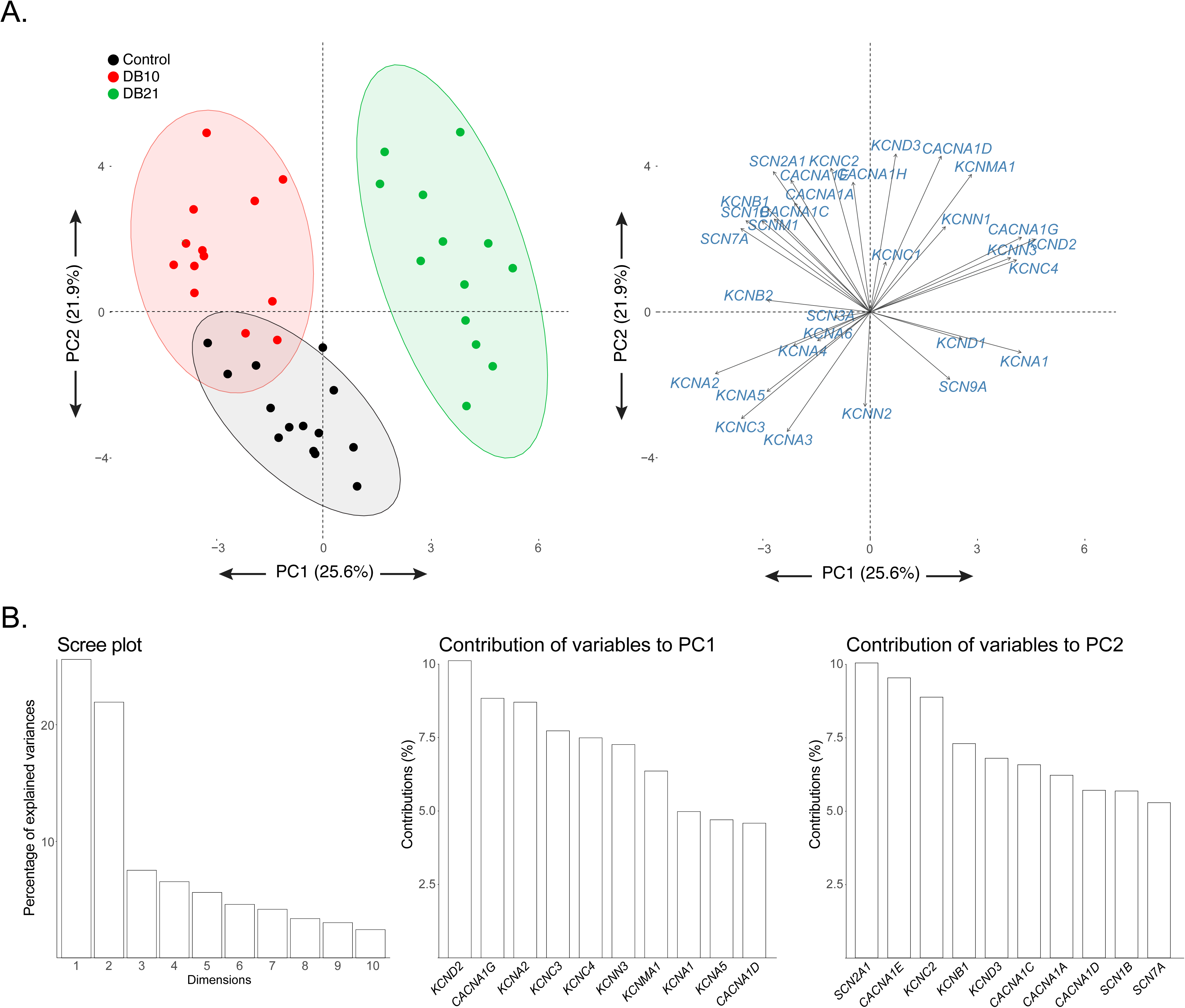
PCA of ion channel mRNA levels across WT, DB10, and DB21 mice. **A.** The first two principal components (PC1 and PC2) define the x-and y-axes respectively. PC1 accounted for 25.6% of the variance while PC2 accounted for 21.9%. The distribution across PC1 and PC2 reveals three distinct ion channel expression profiles for WT, DB10, and DB21 animals. **B.** Scree plot demonstrating the amount of variance accounted for across the first 10 principal components. The majority of the variance is accounted for in the PC1 and PC2, while the remainder contribute far less to the variance in the data. **C.** Post-hoc analysis of the variables that contribute to the variance in PC 1 and PC2 reveals distinct channel subunits that account for the distinct patterns of expression seen in all three experimental groups.

## DISCUSSION

Diabetic cystopathy has formally been documented since 1864 (Faerman et al. 1971), yet the only direct attempt to document how diabetes impacts efferent bladder innervating neurons was by Tompkins et al., (2013) who focused largely on synaptic properties. Therefore, in this study we attempted to identify for the first time the impacts of a type II diabetes model on the intrinsic properties of bladder-innervating parasympathetic neurons in female mice. We hypothesized that passive properties, overall excitability and action potential properties in MPG neurons would change in an age-dependent manner within the diabetic condition. Furthermore, we examined underlying changes in mRNA levels for voltage-dependent ion channels in the whole ganglion as a reflection of the most salient of changes in excitability and firing of MPG neurons.

Lepr ^db/db^ mice on the C57Bl6/J strain used here model hyperglycemia and persistent hyperinsulinemia with beta cell hypertrophy, whereas those on the BKS background strain progress to beta cell failure (Hummel et al. 1972). Although the mice in this study were diabetic at both ages, their hyperglycemia was somewhat worse at 11 weeks than at 21 weeks, consistent with previous observations in this strain (Breyer et al. 2005; Sullivan et al. 2007). This improvement was consistent with the constant, or slightly increasing serum insulin concentrations at 21 weeks, indicating that while the beta cells are not able to adequately compensate for insulin resistance, they have not undergone beta cell failure.

### Diabetes interacts with time to first increase then decrease MPG neuron excitability

One of our most important findings was that excitability of MPG neurons in Lepr^db/db^ females changes nonlinearly with the amount of time exposed to the diabetic environment. Specifically, we observed that 10 week Lepr^db/db^ mice had an increased probability of firing two or more spikes, and decreased rebound spike latency; both of which are measures of increased overall neuronal excitability. In contrast, week 21 Lepr^db/db^ neurons changed in the opposite direction, exhibiting a decrease in spike probability and increased rheobase, both of which are indications of lower overall excitability. This trend of a change in firing properties at 10 weeks that is then rectified at 21 weeks is also seen in the characteristics of action potentials of MPG neurons. Specifically, spikes are narrower (as seen by a decreased half-width, increased ascending slope and enhanced decay slope, and decreased decay time in DB10 animals), and AHPs are shorter are in the DB10 animals but return to control levels in the DB21 group. These data are consistent with the hypothesis that long-term diabetes results in change in MPG neuron that potentially are compensated for via long-term plasticity mechanisms in the DB21 animals. Regardless of whether the changes in excitability and action potential properties are compensatory or represent disease progression in the context of the entire LUT, there is clear change in the characteristics of MPG neurons over the course of 10-21 weeks in diabetic animals.

Because normal bladder function and diabetic cystopathy are the result of a complex interaction of the activities of sympathetic, parasympathetic, and sensory feedback mechanisms combined with bladder compliance and detrusor properties, it is not possible to interpret our results directly in context of mechanisms of changes in bladder output as a result of diabetes. However, the time course of changes in MPG neuron excitability mirror those of bladder output over the progression of diabetic cystopathy in rodents. For example, in type I diabetic mice there is a sharp decline in basal bladder pressure and mean threshold pressure from weeks 9-12 that then is rectified to control levels by 20 weeks (Daneshgari et al. 2006a). Similar time course changes in bladder output features as measured by cystometrogram have been documented in rats as well, whereby diabetic bladders transition from compensated to decompensated states between 9-12 weeks following onset of diabetes (Daneshgari et al. 2006b). Therefore, while a direct mechanistic link to bladder output cannot be made from our data, the results are consistent with the overall change in LUT output over the time course of diabetic cystopathy.

### Potential mechanistic insights via interpretation of changes in mRNA levels

It would be an over-interpretation to directly infer mechanisms underlying physiological changes in single neurons from data collected at the mRNA level from whole ganglia. However, the steady-state mRNA levels for ion channels provide a high-throughput opportunity to generate hypothesis regarding underlying mechanistic changes in ionic currents. Indeed, we see multiple changes in channel mRNAs that are consistent with the physiological changes we report. In this section, we will highlight some of the most salient changes we found at the level of neuronal properties, and cautiously hypothesize on potential underlying mechanism via changes at the mRNA level.

In 10-week Lepr^db/db^ mice we observed that although rheobase did not change, several properties associated with excitability did change. The most salient of these properties was the increase of the probability of firing 2 or more spikes. As shown in Figure 3A and also previously documented (Suzuki and Rogawski 1989; Tompkins et al. 2013), MPG neurons can spike once or many times, a phenomena observed in many other autonomic neuron types (Cassell et al. 1986; Malin and Nerbonne 2001; Springer et al. 2015). One obvious possible mechanism for generating multiple spikes is the activation of a depolarizing current with somewhat slower kinetics that can maintain depolarization above threshold to allow for the generation of multiple spikes. Our data show changes in channel expression that are consistent with this hypothesis. In particular, there is a significant increase in mRNA levels for the calcium channels *CACNA1A, CACNA1E*, and *CACNA1H* in DB10 animals relative to both WT and DB21. Furthermore, *CACNA1E* levels overall are the most abundant calcium channel mRNA that we detected, with levels an order of magnitude higher than any other calcium channel subunit. *CACNA1E* encodes the R-type current Ca_v_2.3, which is known to make up ~25% of total calcium current in rat MPG neurons (Won et al. 2006), and Ca_v_3.2 – encoded by *CACNA1H* – is the predominant T-type channel in rat MPG (Lee et al. 2002). R-type currents are known to influence bursting output in CA1 pyramidal neurons (Metz et al. 2005), and T-type currents in many neurons trigger low-threshold spikes, which in turn generate bursts of action potentials (Perez-Reyes 2003). While the interplay of multiple calcium channel subunits and the remaining ionic conductances of the cell are quite complex in terms of generating different output patterns, increases in R-type and T-type currents would be a feasible way to influence the multiple spiking phenotype in DB10 animals.

We also observed increased ascending and descending spike slopes, and decreased spike half-width in the DB10 animals relative to WT and DB21. We observed increased expression of mRNAs coding for Na_v_1.2 *(SCN2A1)*, as well as a trend for increased Na_v_β1 (SCN1B) expression, in the DB10 animals. Na_v_1.2 channels are largely localized to the axons and initial segments of unmyelinated neurons (Vacher et al. 2008), which is consistent with the post-ganglionic fibers of the MPG. Na_v_β1 is known to interact with Na_v_1.2 channels to increase surface expression of these subunits (Isom et al. 1995). Therefore, increases in expression of *SCN2A1* and *SCNB1* would be expected to increase AP upstroke, which was observed for depolarizing current injections but not rebound spikes. In addition, K_v_3.2 and K_v_2.1 are potassium channels with relatively high threshold activation and fast deactivation such that they have both been implicated as large contributors to the AP repolarization and as a consequence decrease AP width (Rudy and McBain 2001; Liu and Bean 2014). The upregulation of their constituent mRNA subunits *(KCNC2* and *KCNB1* respectively) is entirely consistent with the firing phenotypes observed.

As we observed increases in ascending spike slope to depolarizing current injections, but not rebound current injections in DB10 neurons, it is possible that high threshold potassium currents with fast deactivation time constants could play a role in permitting multiple spikes during depolarizing current injections by enhancing sodium channel de-inactivation. As K_v_2 family members are known to encode delayed rectifiers, and K_v_3 channels have fast activation and deactivation rates associated with sustained higher-frequency firing, then the fact that we see increased Kv3.2 *(KCNC2)* and K_v_2.1 *(KCNB1)* expression is also consistent with these results. While this is well documented for K_v_3.2 as it is a high threshold, fast deactivating (Weiser et al. 1994; Rudy and McBain 2001) ion channel, it is less clear that this is the case for K_v_2.1 as it is intermediate in these parameters. For example, in the superior cervical ganglion (SCG) neurons, K_v_2.1 has somewhat high V_1/2_ activation, slow activation, slow or no inactivation, but a relatively fast deactivation (Liu and Bean 2014). If K_v_2.1 is playing a role in keeping the functional pool of sodium channels available through de-inactivation, as it is thought to do in SCG neurons (Liu and Bean 2014), then we expect its upregulation should prevent decay in spike height during a train of spikes. Therefore, to test this prediction, we examined the ratio of the 2^nd^ to 1^st^ spike height of multi-spiking neurons. A t-test showed that 2^nd^ to 1^st^ spike ratio of DB10 neurons (0.79 ± 0.03, n = 8) were significantly less attenuated than WT neurons (0.48 ± 0.11, n = 2) (t(8) = −4.416, p = 0.002). This suggests that together K_v_3.2 and K_v_2.1 could play a role in keeping sodium currents de-inactivated in the DB10 condition.

DB21 neurons were significantly less excitable than both DB10 and wild type neurons: rheobase was significantly increased, and input resistance was significantly reduced compared to wildtype, and DB21 neurons did not produce a multi-spiking output when stimulated. DB21 neurons had significantly upregulated mRNAs encoding several low threshold slow deactivating potassium currents that would be expected to reduce excitability. The upregulation of *KCNA1* (K_v_1.1) likely contributes to reduced excitability given a relatively low threshold of activation (~-32 mV) and intermediate deactivation time constant (Grissmer et al. 1994). The upregulation of *KCND2* (K_v_4.2) is consistent with the observed decrease in excitability and may explain in part why the DB21 condition did not share the multispiking phenotype with DB10. This is because in cultured rat superior cervical ganglion neurons, Malin and Nerbonne (2000) showed that when K_v_4.2 is overexpressed, the number of neurons displaying the multi-spiking neuron phenotype is decreased, and when the gene is downregulated by expression of a dominant negative transgene, there is a corresponding increase in the multi-spiking phenotype. Interestingly, the authors also report increased input resistance in cells with reduced K_v_4.2, and decreased input resistance with overexpression of K_v_4.2 (Malin and Nerbonne 2001) which is consistent with the significant decrease in R_in_ in DB21 neurons.

## Conclusions

We expected excitability of Lepr^db/db^ neurons to change with increasing intensity from weeks 10 to 21 as diabetes progressed as diabetic neuropathy is a function of both hyperglycemia and duration in humans (Maser et al. 1989; Davies et al. 2006), rats (Mattingly and Fischer 1983; Sasaki et al. 2002) and mice (Giachetti 1978; Hinder et al. 2017; Liu et al. 2017). Contrary to this hypothesis, many properties associated with excitability changed in 10-week animals, but were then resolved closer to wild-type levels in the 21-week animals. These properties include both characteristics of intrinsic excitability (e.g. probability to fire, threshold, and rebound latency) as well as many of the characteristics of the individual action potentials of these neurons as well. Yet, our expression profiling of the MPGs does not simply reveal a resolution at 21-weeks of changes in expression that occur in the 10-week animals. Rather, our PCA reveals that the overall expression patterns in MPGs of wild type, 10-week, and 21-week animals are entirely distinct. Taken together, we suggest that these results represent an apparent compensatory response, where neurons of the MPG in younger diabetic mice are less adapted to hyperglycemia, or the associated bladder cystopathy, than older diabetic mice that have been exposed to hyperinsulinemia for longer times.

## Acknowledgments

This work was funded by a grant from the Missouri Spinal Cord Injuries Research Program (D.J.S.), the Craig H. Neilsen Foundation (D.J.S.), American Diabetes Association Grant 1-14-BS-181 (L.C.S.) and American Heart Association Postdoctoral Fellowship 13POST16910108 (K.A.P.). The authors declare no competing financial interests.

